# Microwave Mutagenesis of Brevibacillus Parabrevis for Enhanced Cellulase Production, and Investigation on Thermostability of this Cellulase

**DOI:** 10.1101/064410

**Authors:** Pinakin Khambhala, Purva Paliwal, Vijay Kothari

**Affiliations:** Institute of Science, Nirma University, Ahmedabad-382481,Gujarat, India

**Author notes:** Corresponding author; Phone: (office).

**Keywords:** Cellulase, Microwave mutagenesis, Thermostability, Fluorescence spectroscopy

## Abstract

Microwave mutagenesis of *Brevibacillus parabrevis* for enhanced cellulase production was attempted. Though microwave treatment could alter the cellulase activity of the test bacterium, none of the mutants obtained were found to be genetically stable, indicating the reversible nature of microwave-induced mutation(s). Thermal stability of the *B. parabrevis* cellulase was also investigated. This enzyme was found to be capable of retaining its activity even after heat treatment (50-121°C, for 30-60 min). Fluorescence spectrum revealed a *red shift* in the emission maxima of the heat-treated enzyme preparations, indicating some structural change upon heating, but no major loss of activity was observed. This enzyme was found to be active over a broad temp range, with 90°C as the optimum temp, which is interesting as the producing organism is a mesophile.

## INTRODUCTION

The part of electromagnetic radiation corresponding to the frequency range of 300 MHz – 300 GHz is known as the microwave (MW) region. Thermal effect of the MW radiation is well established. However, there has been a heated controversy regarding the possible athermal (MW specific electromagnetic effects) effects of the MW radiation (**Trivedi *et al*., 2011**). Literature contains reports indicating presence (**Porcelli *et al.*, 1997**) as well as absence (**Sasaki *et al*., 1995**) of the athermal effects of the MW radiation. Few reports (**Gosai *et al*., 2014**) have also appeared describing the mutagenic potential of MW radiation.

*Brevibacillusparabrevis* is a gram-variable, aerobic, rod shaped **(Logan and Vos, 2009)** bacterium known for its cellulolytic activity (**Singh and Bansal, 2013**).Cellulases are enzymes of high industrial significance, particularly thermostable cellulases are of special interest. Either the cellulolytic thermophilic microbes or the purified thermostable cellulases can be of great use in alcoholic fermentation of lignocellulosic wastes (**Van Maris *et al*, 2006**). The present study aimed at obtaining a cellulase overproducing mutant of *B. parabrevis* using MW radiation as mutagenic agent. Besides this thermostability of the cellulase produced by the wild type *B. parabrevis* was also investigated.

## MATERIAL AND METHODS

### Test Organism

*Brevibacillus parabrevis* (MTCC 2708) was procured from Microbial Type Culture Collection (MTCC) Chandigarh.

#### Microwave Treatment

Bacterial suspension was prepared in sterile normal saline, from an active culture growing on nutrient agar (HiMedia, Mumbai). Inoculum density was adjusted to that of 0.5 McFarland standard i.e., 0.08-0.10 at 625 nm. Test culture (5 mL) in sterile screw capped glass vials (15 mL, Merck) was exposed to MW radiation (90 W; 2450 MHz) in a domestic MW apparatus (Electrolux^®^ EM30EC90SS). MW treatment at 90 W was given for three different time durations viz. 2, 4, and 6 min. Vials inside the MW apparatus were placed in an ice-containing beaker, so as to avoid/minimize any thermal heating. Temperature of the microbial suspension after MW treatment at 90 W did not go beyond 14°C; when 90 W treatment was given for 6 min, temperature of the treated microbial suspension was found to be 13.3 ± 0.5°C. The whole MW treatment was performed in an air-conditioned room. Untreated inoculum was used as control. Before MW treatment all, the inoculum vials (including control) were put in ice for 5 min to nullify any variations in initial temperature. Test organism was immediately (in less than 5 min) inoculated onto the medium for screening of cellulolytic potential (**Gupta *et al*, 2012**), following MW treatment. This growth medium contained 0.5 g/L KH2PO4 (Merck, Mumbai), 0.25 g/L MgSO4.7H2O (Merck), 0.2 g/L congo red (S.d. fine-chemi, Mumbai), 2 g/L cellulose (S.d. fine-chemi), 2 g/L gelatin, 15 g/L agar-agar (HiMedia, Mumbai), pH 6.8-7.2.Incubation was made at 35°C for 24 h.

### Estimation of cellulase activity

*B. parabrevis* was grown in a CMC (carboxymethyl cellulose; Merck) supplemented broth (Peptone 0.5 g/L; NaCl 0.5 g/L; Beef extract 0.5 g/L; Yeast extract 0.5 g/L; CMC-Na 20 g/L). Incubation of the experimental tubes was carried out at 35°C for 65 h, under shaking condition (80 rpm). The cell free supernatant obtained after centrifugation of the culture broth (10,000 rpm; 9,390 g for 15 min) was used as crude cellulase preparation. 0.5 mL of the supernatant was mixed with 0.5 mL of 1% CMC, followed by incubation at 50°C for 30 min. The amount of glucose released as a result of cellulase activity was quantified using DNSA colorimetric assay. The international units (IU) of the cellulase was calculated as: IU= (μg of glucose)/ 180 × 30 × 0.5 (**Nigam and Ayyagari, 2008**).

### Screening for mutants

Following the MW treatment of *B. parabrevis* suspension, the treated inoculum was streaked on the screening medium plate (150 mm) containing congo red, and incubated at 35°C for 42-45 h. After the incubation 3 colonies from each plate corresponding to different MW treatments were picked (selection of the colonies was made based on the diameter of the zone of cellulose hydrolysis surrounding the colony), and each colony (a separate code was given to each picked colony) was streaked on to a separate nutrient agar (supplemented with 0.5% CMC) plate. Daughter populations thus generated (after 42-45 h incubation at 35°C) from a single parent colony were then inoculated into the liquid medium described in the preceding section for estimation of enzyme activity, followed by incubation at 35°C for 65 h under shaking condition (80 rpm). After incubation, cellulase activity was estimated for all the MW treated inoculums. Then the plates corresponding to the MW treatment yielding higher cellulase activity were selected for further experiments. Subculturing was done from the plates of cellulase overproducing mutant(s), and daughter population resulting from each subculturing was checked for its cellulase activity (in comparison to the wild type), up to 5 generations.

### Partial purification of the enzyme preparation

*B. parabrevis* was inoculated into CMC containing broth, in 500 mL flasks. Volume of the growth medium in the flask was kept 250 mL. Incubation was carried out at 35°C under shaking condition (80 rpm) for 65 h. Cell free supernatant obtained after centrifugation of the *B. parabrevis* culture broth was subjected to ammonium sulphate (CDH, New Delhi) precipitation, wherein ammonium sulphate was added at the rate of 690 g/L. While adding ammonium sulphate [(NH_4_)_2_SO_4_], the vessel containing the supernatant was kept surrounded by ice, and following that a 20 min incubation inside refrigerator was carried out. This was followed by centrifugation at 7,500 rpm for 45 min at 4°C. The resulting pellet containing the precipitated protein content was dissolved in 25 mM phosphate buffer. This solution was subjected to dialysis (11 KD pore size, 33 mm wide; Sigma Aldrich), wherein the dialysis bag was kept suspended in 25mM phosphate buffer for 12 h under refrigeration. At the end of dialysis the contain inside bag was centrifuged at 10,000 rpm (15 min; 4°C), and the resulting supernatant was used as partially purified cellulase preparation.

### Finding the effect of temperature on enzyme-substrate interaction

0.5 mL of the partially purified enzyme was mixed with 0.5 mL of 1% CMC, followed by incubation for 30 min at different temp in the range 40-100°C. The amount of glucose released as a result of cellulase activity was quantified using DNSA colorimetric assay, and from that the international units (IU) of the cellulase was calculated. Appropriate controls containing only substrate (with no enzyme), and only enzyme (with no substrate) were also included in the experiment.

### Investigating thermostability of the enzyme

The partially purified enzyme was subjected to heating in water bath for 1 h at different temp viz. 50°C, 80°C, and 100°C. Besides this it was also subjected to autoclaving for 30 min. One batch of these heated enzyme preparations was immediately subjected to estimation of cellulase activity. Another batch from the same lot was allowed to cool and undergo renaturation for 2 h. Following renaturation this was also subjected to estimation of cellulase activity. Additionally the renatured sample was also subjected to flurometric analysis to find out whether the reantured sample has undergone any structural change as compared to the unheated control. Fluorescence spectroscopy experiments were performed using cary eclipse fluorescence spectrophotometer, Agilent technology (US). Excitation wavelength used was 280 nm, and the emission was recorded in the range 300-400 nm.

#### Statistical analysis

All the experiments were performed in triplicate, and measurements are reported as mean ± standard deviation (SD). Statistical significance of the data was evaluated by applying *t*-test using Microsoft Excel^®^. *P* values less than 0.05 were considered to be statistically significant.

## RESULTS AND DISCUSSION

### Microwave mutagenesis

Following MW treatment of the *B. parabrevis* inoculum, it was streaked on to congo red containing screening medium, and the colonies surrounded by zone of cellulose hydrolysis bigger than that of control (Table 1) were selected for further experimentation. Each of the nine colonies shown in Table 1 was streaked on to a separate nutrient agar (supplemented with 0.5% CMC) plate. Daughter populations thus generated (after 24 h incubation at 35°C) from a single parent colony were then inoculated into the liquid medium described in the section for estimation of enzyme activity, followed by incubation at 35°C for 65 h under shaking condition (80 rpm). After incubation, cellulase activity was estimated for all the MW treated inoculums (Table 2). Then the plates corresponding to the MW treatment yielding higher cellulase activity were selected for further experiments. Subculturing was done from the plates of cellulase overproducing mutants (i.e. 4A, 6A, 6B, and 6C), and daughter population resulting from each subculturing was checked for its cellulase activity (in comparison to the wild type), till any significant higher cellulase activity was observed (Table 3). Out of the four mutant strain selected, none could maintain the trait of cellulase overproduction after third subculturing. Still we continued the experiments till fifth subculturing, and found all the mutants to revert back to the parent phenotype with respect to cellulase activity.

**Table 1.**
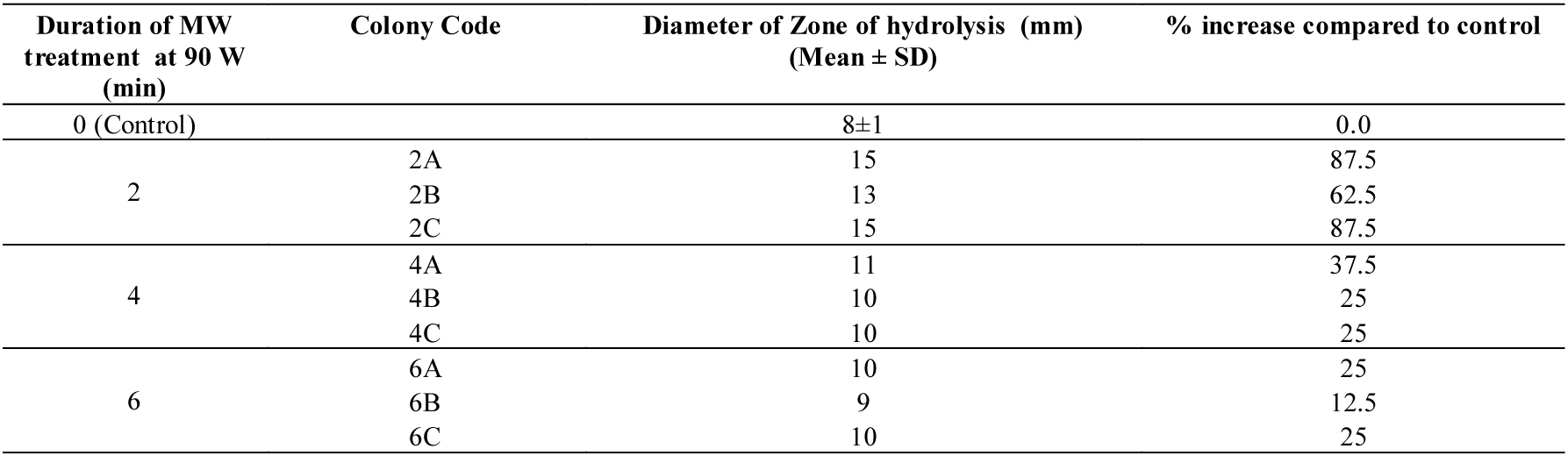
Results of qualitative screening of the cellulolytic potential on congo red medium.

**Table 2.** Quantification of cellulase activity of the population generated from selected colonies.

| Colony Code | Cell density at harvest (OD<sub>660</sub>) (Mean ± SD) | % Change compared to control | Cellulase activity (IU/mL) | % Change compared to control |
| --- | --- | --- | --- | --- |
| Control | 0.716±0.04 | 0.00 | 0.020 | 0.00 |
| 2A | 0.613±0.06 | -14.38 | 0.017 | -15 |
| 2B | 0.614±0.01 | -14.24 | 0.018 | -10 |
| 2C | 0.623±0.01 | -12.98* | 0.023 | 15 |
| 4A | 0.729±0.05 | 1.81 | 0.050 | 150** |
| 4B | 0.577±0.01 | -19.41* | 0.021 | 5 |
| 4C | 0.695±0.00 | -2.93 | 0.023 | 15 |
| 6A | 0.749±0.06 | 4.60 | 0.039 | 95** |
| 6B | 0.737±0.02 | 2.93 | 0.047 | 135** |
| 6C | 0.693±0.00 | -3.24 | 0.049 | 145** |

$p < 0.05$ ; \*\* $p < 0.01$ ; minus sign indicates a decrease over control

**Table 3.** Cellulase activity over multiple subculturings of selected overproducing mutants of *B. parabrevis.*

| Colony designation | Cell density at harvest (OD<sub>660</sub>) (Mean ± SD) | % Change compared to control | Cellulase activity (IU/ ml) | % Change compared to control |
| --- | --- | --- | --- | --- |
| 1<sup>st</sup> generation |  |  |  |  |
| Control | 0.71±0.04 | 0.00 | 0.020 | 0.00 |
| 4A | 0.72±0.05 | 1.81 | 0.050 | 150** |
| 6A | 0.74±0.06 | 4.60 | 0.039 | 95** |
| 6B | 0.73±0.02 | 2.93 | 0.047 | 135** |
| 6C | 0.69±0.00 | -3.24 | 0.049 | 145** |
| 2<sup>nd</sup> generation |  |  |  |  |
| Control | 0.37±0.00 | 0.00 | 0.029 | 0.00 |
| 4A | 0.42±0.00 | 12.5** | 0.026 | -10.34 |
| 6A | 0.47±0.01 | 26.32** | 0.045 | 55.17 |
| 6B | 0.34±0.02 | -9.30* | 0.043 | 48.27* |
| 6C | 0.48±0.00 | 30.05** | 0.036 | 27.58* |
| 3<sup>rd</sup> generation |  |  |  |  |
| Control | 0.66±0.00 | 0.00 | 0.030 | 0.00 |
| 4A | 0.53±0.07 | -19.88 | 0.042 | 40** |
| 6A | 0.61±0.00 | -7.62** | 0.042 | 40** |
| 6B | 0.52±0.02 | -24.89** | 0.039 | 30 |
| 6C | 0.52±0.00 | -24.74** | 0.045 | 50 |
| 4<sup>th</sup> generation |  |  |  |  |
| Control | 0.71±0.00 | 0.00 | 0.015 | 0.00 |
| 4A | 0.76±0.02 | 7.11 | 0.017 | 13.33 |
| 6A | 0.68±0.02 | -4.18 | 0.023 | 53.33 |
| 6B | 0.68±0.02 | -4.04 | 0.022 | 46.66 |
| 6C | 0.71±0.01 | -0.13 | 0.020 | 33.33 |
| 5<sup>th</sup> generation |  |  |  |  |
| Control | 0.52±0.00 | 0.00 | 0.021 | 0.00 |
| 4A | 0.49±0.02 | -5.71 | 0.022 | 4.76 |
| 6A | 0.50±0.02 | -3.23 | 0.022 | 4.76 |
| 6B | 0.48±0.00 | -7.61 | 0.026 | 23.80 |
| 6C | 0.50±0.02 | -3.61 | 0.026 | 23.80 |

\* $p < 0.05$ ; \*\* $p < 0.01$ ; minus sign indicates a decrease over control.

Though the MW radiation could exert its mutagenic effect on *B. parabrevis* cellulase activity, the mutants obtained were not found to be genetically stable. It can be said that the MW induced mutations in the cellulase synthesizing/secretion machinery observed in the study were of reversible nature. Literature contains reports indicating the MW induced mutations to be stable, as well as those indicating otherwise. **Pasiuga *et.al*. (2007)** reported disappearance of low-level MW induced effects after few generations in *Drosophilla malenogaster*. **Kothari *et.al.* (2014)** also showed mutagenic effect of MW radiation on exopolysaccharide production in *Xanthomonas campestris* to be of reversible type.

Whatever alterations in growth and cellulase activity of MW exposed B. parabrevis were observed in the study, seem largely owing to the non-thermal effects of MW radiation, as MW treatment in the study was provided at relatively low power (90 W) and the temperature of the MW treated suspension did not go beyond 14°C. Much cannot be said with certainty regarding the mode of action of MW radiation on living systems **Bollet *et.al.* (1991)** Reported alteration in the cell membrane permeability owing to MW treatment, which in part may contribute to the non-thermal effects of MW on microbial cells. Besides thermal and non-thermal effects, micro-thermal effects incorporating ‘undetectable’ thermal mechanism may also be responsible for the biological effects of microwave radiation (**Shamis *et al*, 2012**).

### Effect of temperature on the activity of *B. Parabrevis* cellulase

#### Finding the effect of temperature on enzyme-substrate interaction

This experiment was done in two sets. For the first set organism was grown in a medium containing 0.5% cellulose as the major carbon source, and Whatman paper discs (5 discs of diameter 5 mm, amounting to 10.68±0.16 mg, in each experimental tube) were used as the substrate during enzyme assay. In the second set, organism was grown in a medium containing 2% CMC as the major carbon source, and 1% CMC was used as the substrate during enzyme assay. Temperature at which an incubation for 30 min was made allowing the enzyme to work on substrate ranged from 40-100°C. Cellulase activity obtained at different temp for both set of experiments is shown in Table 4 and Figure 1. In the first experimental set the cellulase activity was found to be maximum at 90°C. Enzyme activity at 40°C was statistically equivalent to that of 90°C. At all other temperatures the enzyme activity was found to be lesser (30-70% of that found at 90°C). In the second set of experiments, maximum enzyme activity was observed at 90°C, however the cellulase activity in the whole range of 40-100°C was almost similar. Though statistically significant but minor decrease in activity was obtained at 50°C and 60°C. This experiment was also performed with unpurified crude enzyme (i.e. the cell free supernatant used directly as an enzyme source), and maximum activity was observed at 90°C (data not shown). The level of cellulase activity in second set (using CMC as the substrate for enzyme assay) of experiments is almost double than that observed in the first set (using Whatman paper as the substrate for enzyme assay). This may possibly be due to different affinities of the *B. parabrevis* cellulase for two different substrates. Many factors contribute in determining how challenging cellulase activity assays can be. Few among these factors are the degree of homogeneity and purity of the cellulase sample; solubility of the substrate; the complicated relationship between physical heterogeneity of the cellulosic materials, and the complexity of cellulase enzyme systems (synergy and/or competition)(**Zhang *et al*., 2015**).

**Figure 1.**
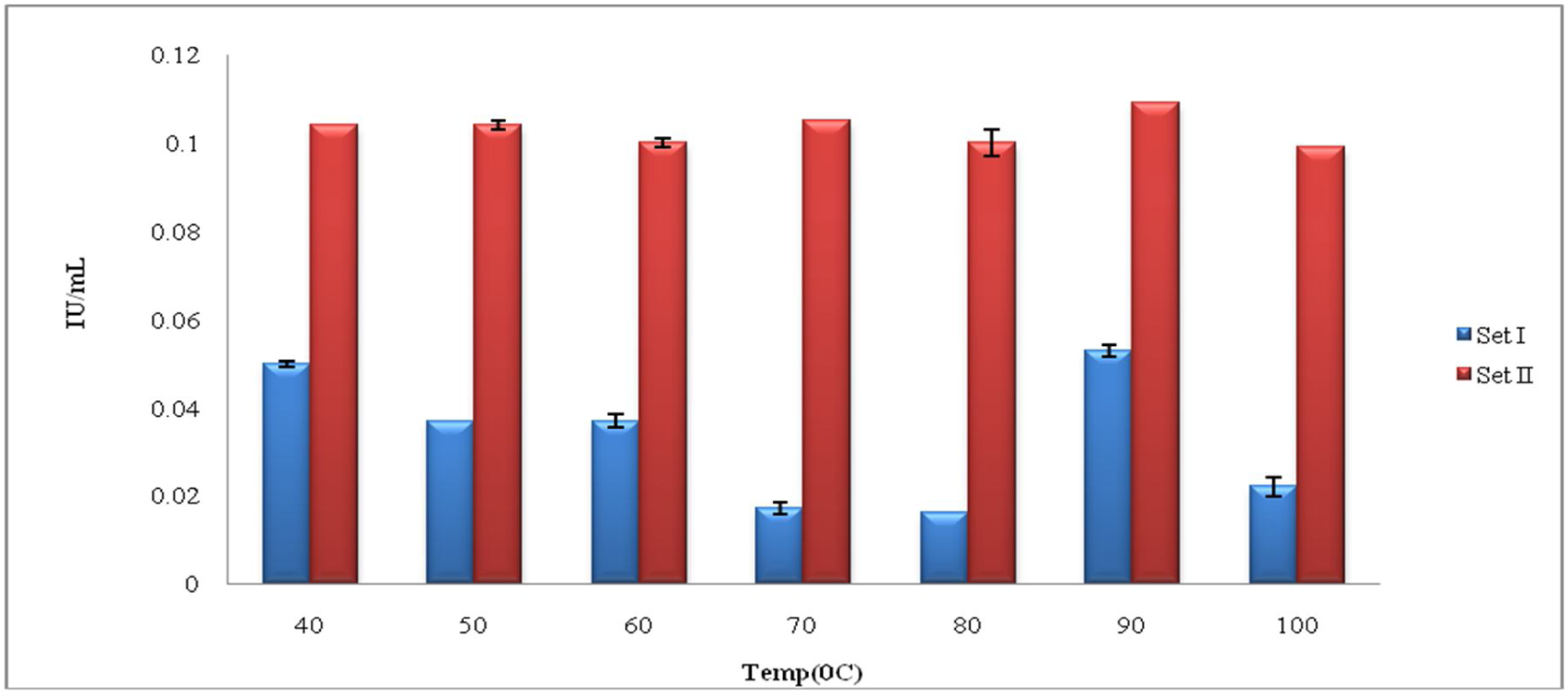
Cellulase activity as a function of temperature. **Set I** Cellulose as growth substrate, and Whatman paper as substrate during enzyme assay. **Set II** CMC as growth substrate, as well as substrate during enzyme assay

**Table 4.**
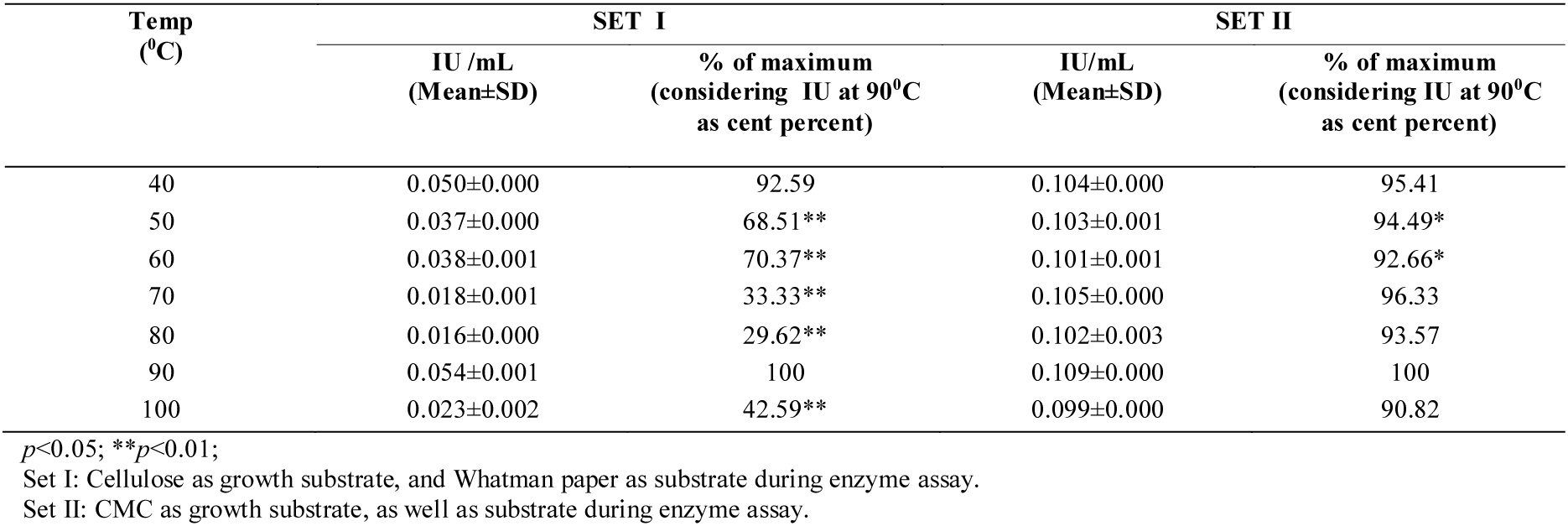
Cellulase activity as a function of temperature.

| Temp<br>(°C) | SET I |  | SET II |  |
| --- | --- | --- | --- | --- |
|  | IU/mL<br>(Mean±SD) | % of maximum<br>(considering IU at 90°C<br>as cent percent) | IU/mL<br>(Mean±SD) | % of maximum<br>(considering IU at 90°C<br>as cent percent) |
| 40 | 0.050±0.000 | 92.59 | 0.104±0.000 | 95.41 |
| 50 | 0.037±0.000 | 68.51** | 0.103±0.001 | 94.49* |
| 60 | 0.038±0.001 | 70.37** | 0.101±0.001 | 92.66* |
| 70 | 0.018±0.001 | 33.33** | 0.105±0.000 | 96.33 |
| 80 | 0.016±0.000 | 29.62** | 0.102±0.003 | 93.57 |
| 90 | 0.054±0.001 | 100 | 0.109±0.000 | 100 |
| 100 | 0.023±0.002 | 42.59** | 0.099±0.000 | 90.82 |

$p < 0.05$ ; \*\* $p < 0.01$ ;
Set I: Cellulose as growth substrate, and Whatman paper as substrate during enzyme assay.
Set II: CMC as growth substrate, as well as substrate during enzyme assay.

Cellulases capable of acting on their substrate at high temperatures are of interest for various industrial applications, as process is employing cellulases at higher temperature are less prone to contamination. Cellulolytic activity at high temperature is of special interest with respect to alcohol production from lignocellulosic wastes (**Van Maris *et al*., 2006**). In the present study, we have found the cellulase activity to be optimum at a temperature as high as 90°C, this is particularly interesting considering that the cellulase producing organism used here is mesophilic with an optimum growth temperature of 35°C. This will also be interesting to investigate why a mesophilic bacterium need to produce a thermotolerant enzyme. Genes coding for such enzymes may also be useful while practicing genetic engineering for construction of other superior cellulase producing strains. Many microorganisms are known for their good cellulolytic potential, but their cellulases are functioning optimally at mesophilic temperatures. For example, **Li *et.al***. (**2010**) have shown optimum temperature CMCase activity from *Trichodermaviride* to be 50°C.

#### Investigating thermostability of the enzyme

After observing that the optimum temperature of the *B. parabrevis* cellulase is on the higher side, we proceeded for investigating whether this enzyme undergoes any structural modification after being subjected to heating. These experiments were performed in three different sets. Set I employed cellulose as the growth substrate, and Whatman paper as as the substrate during enzyme assay. Set II employed CMC as the growth substrate as well as substrate during enzyme assay. Set III employed CMC as the growth substrate, and Whatman paper as the substrate during enzyme assay. Here, we subjected the partially purified enzyme solution to heating at 50°C, 80°C, 100°C for 1 h, and 121°C (autoclaving) for 30 min. Results of the cellulase activity assay for the enzyme preparation(s) immediately after the heat treatment, and following cooling are presented in Table 5-7. Fluorescence spectrum of these cooled enzyme samples (for set I and II) were also generated.

**Table 5.**
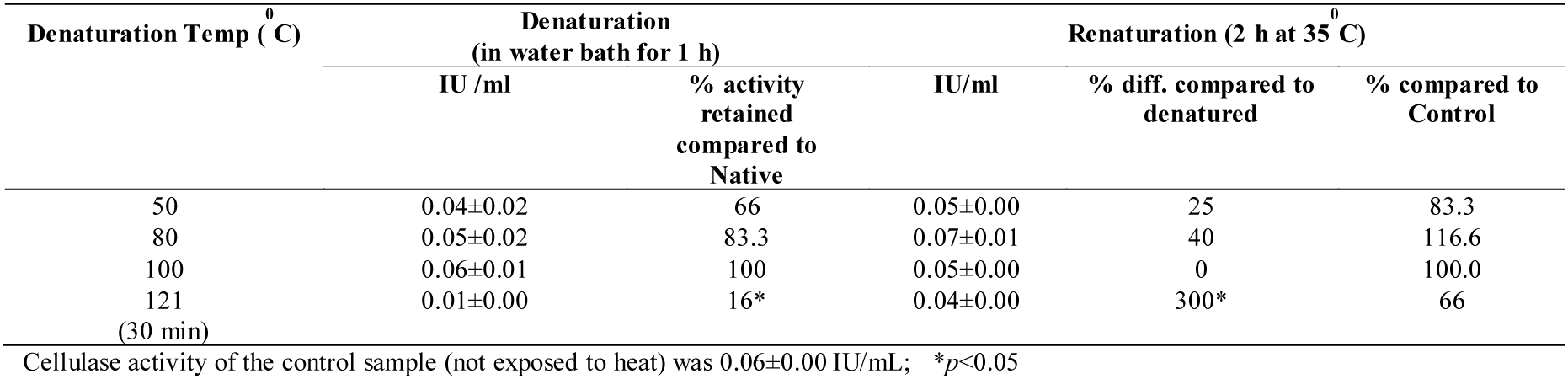
Cellulase activity of the heat denatured, and renatured enzyme preparation (set I).

**Table 6.** Cellulase activity of the heat denatured, and renatured enzyme preparation (set II).

| Denaturation Temp (°C) | Denaturation<br>(in water bath for 1 h) |  | Renaturation (2 h at 35°C) |  |  |
| --- | --- | --- | --- | --- | --- |
|  | IU /ml | % activity retained compared to Native | IU/ml | % diff. compared to denatured | % compared to Control |
| 50 | 0.099±0.001 | 90.00* | 0.104±0.002 | 5.05 | 94.54 |
| 80 | 0.096±0.001 | 87.27* | 0.100±0.003 | 4.16 | 90.90 |
| 100 | 0.098±0.001 | 89.09* | 0.101±0.002 | 3.06 | 91.81 |
| 121<br>(30 min) | 0.096±0.000 | 87.27* | 0.100±0.002 | 4.16 | 90.90 |

Cellulase activity of the control sample (not exposed to heat) was 0.110±0.002IU/mL; \* $p$ <0.05.

**Table 7.** Cellulase activity of the heat denatured, and reantured enzyme preparation (set III).

| Denaturation Temp (°C) | Denaturation<br>(in water bath for 1 h) |  | Renaturation (2 h at 35°C) |  |  |
| --- | --- | --- | --- | --- | --- |
|  | IU /ml | % activity retained compared to control | IU/ml | % diff compared to denatured | % compared to control |
| 50 | 0.223±0.000 | 83.52** | 0.249±0.04 | 111.65* | 93.25 |
| 80 | 0.229±0.007 | 85.76* | 0.219±0.012 | 95.63 | 82.02* |
| 100 | 0.182±0.012 | 68.16* | 0.202±0.025 | 110.98 | 75.65 |
| 121 (30 min) | 0.237±0.015 | 88.76 | 0.229±0.001 | 96.62 | 85.76* |

Cellulase activity of the control sample (not exposed to heat) was 0.267±0.005 IU/mL; \* $p$ <0.05; \*\* $p$ <0.01

During the experiments of set I, enzyme activity in the samples heated at 50-100°C was significantly not different than that in unheated control. However the autoclaved enzyme preparation suffered a loss of 83.34% activity compared to the control; this sample regained activity equivalent to the control after cooling. Thus except for autoclaving, the enzyme activity was not affected by heating. Though the activity of all the heat treated enzyme samples (after cooling) was at par to that of unheated control, structures of the reantured enzyme samples did not seem to be the same, as indicated by a shift in their fluorescence spectrum. Fluorescence spectrum of the native (unheated control) enzyme preparation exhibited an emission maxima at 341 nm, whereas this value was higher for the renatured enzyme samples (which were previously subjected to heat treatment) indicating a red shift (Fig. 2). For the experiments of set II, there was a minor (11-13%) decrease in the enzyme activity measured immediately after heat treatment; however following renaturation upon cooling the experimental enzyme preparations showed no significant difference with respect to their activity compared to the control. Fluorescence spectrum obtained in this set of experiments (spectrum not shown to avoid redundancy) was similar to that obtained for set I, exhibiting a red shift in the emission maxima of the heat treated enzyme preparations. For the third set of experiments, enzyme activity after heat treatment suffered maximum (32%) decrease at 100°C, however following cooling it regained the activity equivalent to that of control.

**Figure 2.**
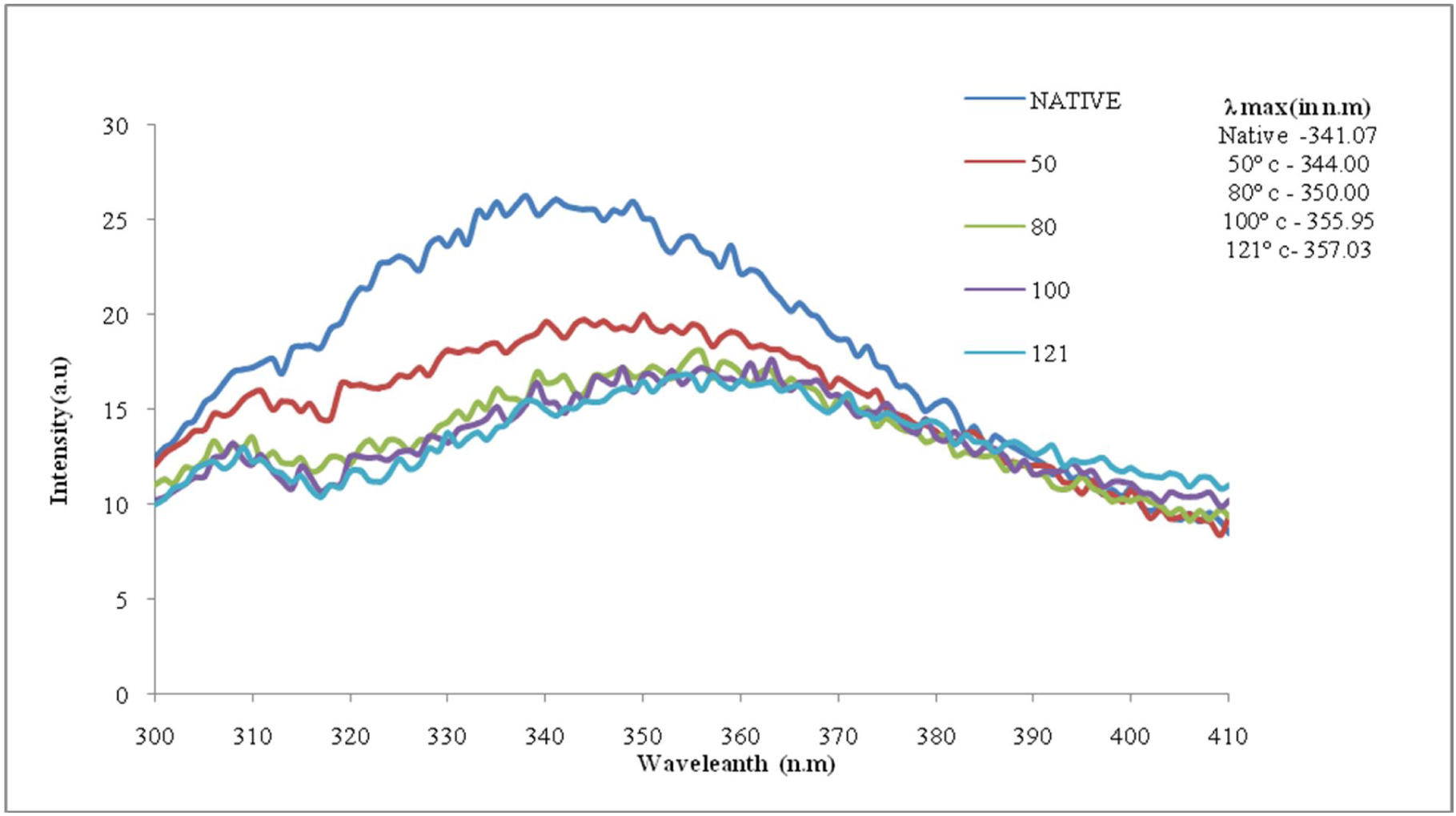
Structure of enzyme preparation after renaturation (Set I).

From the above described experiments regarding thermostability of the *B. parabrevis* cellulase, it is evident that this enzyme does not show any major significant reduction in its activity when measured immediately after heat treatment (i.e. retains considerable activity even after heat treatment), and upon renaturation (after cooling) regains activity almost at par to that of native (unheated control). However the structure of this enzyme does undergo some change(s) upon heat treatment, as indicated by the fluorescence spectrum. But this altered structure also exhibits catalytic efficiency almost equivalent to that of native structure. It may be assume that the active site of this cellulase is not getting distorted even after structural alteration(s) owing to heat treatment, neither its accessibility to the substrate is reduced much. In our study the fluorescence spectrum of the heat treated enzyme preparation exhibited an increase in the emission maxima as compared to that of native, and the magnitude of this shift increased with increase in temp (Fig 2). This phenomenon of red shift can be considered as an indication of unfolding and denaturation of the enzyme structure, and revelation of more hydrophobic amino acid to the surface. The indole group of tryptophan is the dominant fluorophore in proteins, which absorbs near 280 nm, and emits near 340 nm. The emission of indole may shift to longer wavelengths (red shift) when the protein is unfolded (**Joseph *et al.*, 2006**). The indole group of tryptophan residues in proteins is a solvent-sensitive fluorophore, and the emission spectra of indole can reveal the tryptophan residues in proteins. The emission from an exposed surface residue is known to occur at longer wavelengths than that from a tryptophan residue in the protein’s interior. This phenomenon apparent in Figure 2 is characterized by a shift in the spectrum of a tryptophan residue upon unfolding of a protein, and the subsequent exposure of the tryptophan residues to the aqueous phase. Before this unfolding, the residue is likely to be shielded from the solvent by the folded protein. Such unfolding of a cellulase may not always affect its substrate binding properties. Binding behavior of *Trichoderma reesei* celullases was shown not to be adversely affecting at temperature above 50^0^C (**Andreaus *et al.*, 1999**). Tryptophan residue has been indicated as key amino acid in the structure of *T. reesei* cellulase (**Nakamura *et al.*, 2013**), as well as in cellulase of Bacillus species (**Ozaki *et al.*, 1991**). An increased frequency of exposed aromatic residues is believed to underline a stabilizing effect in enhancing thermal stability of proteins (**Chakravarty *et al.*, 2002**).

## CONCLUSION

The study attempted at MW mutagenesis of *B. parabrevis* for enhanced production of thermostable cellulase. Though none of the four overproducing mutant lines obtained was found to be genetically stable, our study positively indicate the possible contribution of athermal MW effects towards their mutagenic potential. Investigation on the thermostability of the cellulase activity (performed with the wild type) revealed that the enzyme activity is not destroyed much even after heat treatment, despite certain structural changes (revealed from fluorescence spectrum) taking place owing to heat treatment. This enzyme was found to be catalytically active over a broad range of temperature. To the best of our awareness, this is the first description regarding thermostability of the *B. parabrevis* cellulase. More detailed investigation on this cellulase is warranted to understand the molecular basis underlining the thermostable nature of this protein. Cellulases capable of working at high temperature and retaining activity even after considerable heat exposure can find many interesting industrial applications **(Haki *et al*., 2003**). It is noteworthy that this thermostable cellulase is not from a thermophilic organism, but from a mesophile. Mesophiles are easier to handle in lab as well as during large scale fermentative processes.

## Acknowledgments

Authors thanks Nirma Education and Research Foundation (NERF) for financial and infrastructural support; Mili Das and her students (Palak Patel and Krupali Parmar) for their help in fluorescence spectroscopy.

